# Physiological Responses to Caprine Coccidiosis During a 100-year Rain Event in Oklahoma: An Opportunistic Case Study with a implications for climate change and clinical diagnosis

**DOI:** 10.1101/2022.01.25.477737

**Authors:** Malcolm L. McCallum, Yonathan Tilahun, Jessica Quijada Pinango, Zaisen Wang

## Abstract

The interaction between inclement weather and disease acquisition is a long-recognized relationship. In the case of coccidiosis, a parasitic infestation of the intestines, wet weather is known to promote incidences in livestock. Our opportunistic investigation tracks blood chemistries of goats that were exposed to coccidian in a 100 year rain event in Oklahoma. Our results suggest a potentially patterned response of blood chemistries that may be developed into a clinical tool in the future and provide warnings for producers of the risks associated with growing incidences of excessively wet weather related to climate change.

## Introduction

Coccidiosis is an important contagious syndrome that afflicts virtually all vertebrate organisms, including livestock (Jolley and Bardsley 2006; Foreyt 1990). Several species of *Eimeria* afflict goats, with the most pathogenic species being *E. arloingi*, *E christenseni*, and *E. ovinoidalis* (ConsTable 3020; Chartier and Paraud 2012; Aumont et al. 1984). These parasites are intracellular and result in epithelial sloughing, inflammatory response, blood loss, and potentially death (ConsTable 3020; Tafti and Mansourian 2008; Valentine et al. 2007).

We explored if clinical blood chemistry parameters responded to coccidia exposure resulting from wet weather and if this could be modelled to create a predictive tool for early diagnosis of exposure to coccidians. Such a tool could serve as a proactive measure to reduce parasitic impacts on dairy goats. We hypothesized that days spent outside should reflect exposure to coccidia and ultimately parasite loads. So, some hematological parameters maybe predictive of the level of exposure animals experienced. We surmised that some parameters should respond to coccidia loads and be useful for predictive regression modeling to detect infestations before they become outwardly symptomatic.

## Materials and Methods

This study took place at the North American Institute for Goat Research of Langston University (Langston, Oklahoma, USA). Rainfall during this period was 337% above normal making it the wettest May in history (Collington 2019) and resulted in declaration of four separate national disasters for storms and flooding (FEMA EM-3411 [24 May], DR-4438 [31 May], DR-4453 [11 July], DR4456 [6 August]). Hence, the potential for pasture-housed goats to develop coccidiosis was extreme, as indicated in earlier studies (e.g., O’Callagham 1989; Anene et al. 1994; Abo-Shehada and Abo-Farieha 2003).

Fifty-five 3-month-old Alpine bucklings (BW = 5.54, SE = 0.201) were randomly assigned to pens and in elevated pens (1.2 x 1.2 m) and believed to be nematode-free without signs of coccidiosis. The amount of time each of these animals had spent on pasture was recorded. They were fed a moderate-quality oat hay (12% protein) ad-libitum (about 1% of body weight (BW) on a dry matter (DM) basis). They also had continuous access to a mineral block, water. This was supplemented with our Langston Moderate Protein Pellet (Hart and Goetsch 2001; Table 1) initially fed at 250 g per head and increasing until daily intake was capped at 500 g/kid/day. Goats were divided into five different treatments for an unrelated study on Barber Pole Worms, *Haemonclues contortus* (*Hc*). It involved a control group (n = 15), and four treatment groups (n = 10 goats/treatment) receiving oral doses of 2500 – 10,000 infective L3 *Hc* larvae obtained from Scott Bowdridge (West Virginia University) at the beginning of the study. Blood and fecal samples were collected from each animal before oral applications and then afterwards on days 14, 28, and 42 post-artificial *H.c*. infestation. Blood was sampled with EDTA tubes via the jugular vein. Blood samples were sent to the Oklahoma Animal Disease Diagnostic Laboratory (Stillwater, OK; Accession No. DLBC74368722) for hematological and blood chemistry analysis of 38 different parameters (Table 2). Although fecal samples were screened for Oocysts using McMaster Method (MAFF 1986; performed by JQ) to manage coccidiosis throughout the study, these data were not collected in a sufficiently regimented fashion required for statistical analysis. Hence, time on pasture (exposure) was used as a surrogate for parasite load as previous work has demonstrated that goats pastured during rainy seasons have significantly more coccidian than those pastured at other times (Gwaze et al. 2010).

**Table 1.** Calculated composition on a dry matter basis and ingredients for the Langston Moderate Protein Pellet diet (Hart and Goetsch 2001).

| Component |  | Ingredient | %<br>Air dry |
| --- | --- | --- | --- |
| Crude Protein (% DM) | 20.30 | Cottonseed hulls | 15.00 |
| Total Digestible Nutrients (% DM) | 74.10 | Alfalfa meal | 20.00 |
| Calcium (% DM) | 1.00 | Ground corn | 13.40 |
| Phosphorus (% DM) | 0.50 | Soybean meal | 25.00 |
| Copper ppm | 16.30 | Pellet partner | 18.85 |
| Selenium ppm | 0.32 | Dicalcium carbonate | 5.00 |
| Zinc ppm | 142.00 | White salt | 1.10 |
| Fe, minimum % air dry | 3.55 | Ammonium chloride | 0.50 |
| Zn, minimum % air dry | 3.77 | Trace mineral and |  |
| Mn, minimum % air dry | 1.90 | vitamin premix | 0.25 |
| S, minimum % air dry | 1.20 |  |  |
| K, minimum % air dry | 1.07 | **Component (cont.)** |  |
| Mg, minimum % air dry | 0.79 | d-Pantothenic acid, min. mg/pound | 1,840 |
| I, minimum % air dry | 0.046 | Niacin, Min. mg/pound | 5,000 |
| Co, minimum % air dry | 0.009 | Vitamin B-6, min mg/pound | 14 |
| Se, minimum % air dry | 0.002 | Folic Acid, min mg/pound | 20 |
| Vitamin A, Min. I units/pound | 1,200,000 | Choline, min. mg/pound | 21,620 |
| Vitamin D<sub>3</sub>, Min. I units/pound | 400,000 | Biotin, min. mg/pound | 1 |
| Vitamin E, Min. I units/pound | 1,500 | Methionine, Minimum % | 4.95 |
| Vitamin B-12, min. mg/pound | 5.00 |  |  |
| Vitamin K, min. mg/pound | 165 |  |  |
| Riboflavin, min. mg/pound | 1,100 |  |  |

**Table 2.** Results of General Linear Model for each set of predictors and hematological responses in Alpine Dairy Goats. (Norm = data was normalized; **Bold** = significant and marginally significant for predictors of the number of days outside [i.e., exposure to coccidia])

| Response | Units | Ref. Range | Predictor | df<sub>1</sub> | df<sub>2</sub> | F | P |
| --- | --- | --- | --- | --- | --- | --- | --- |
| PCV | % | -- | Sample Date | 2 | 95 | 10.29 | < 0.001 |
| PCV | % | -- | Days outside | 2 | 95 | 1.22 | 0.301 |
| Total Protein | g/dL | 6.5 – 8.8 | Sample Date | 2 | 95 | 4.39 | 0.015 |
| **Total Protein** | **g/dL** | **6.5 – 8.8** | **Days outside** | **2** | **95** | **7.63** | **0.001** |
| Albumin (Norm) | g/dL | 3.0 – 4.0 | Sample Date | 2 | 95 | 17.94 | < 0.001 |
| Albumin (Norm) | g/dL | 3.0 – 4.0 | Days outside | 2 | 95 | 0.46 | 0.634 |
| Globulin | g/dL | 2.5 – 5.8 | Sample Date | 2 | 95 | 4.33 | 0.016 |
| **Globulin** | **g/dL** | **2.5 – 5.8** | **Days outside** | **2** | **95** | **7.11** | **0.001** |
| A/G ratio (Norm) | -- | 0.53 – 1.77 | Sample Date | 2 | 95 | 10.49 | < 0.001 |
| **A/G ratio (Norm)** | **--** | **0.53 – 1.77** | **Days outside** | **2** | **95** | **4.36** | **0.015** |
| AST (Norm) | IU/L | 60 – 290 | Sample Date | 2 | 95 | 1.85 | 0.163 |
| AST (Norm) | IU/L | 60 – 290 | Days outside | 2 | 95 | 1.74 | 0.181 |
| ALT(SGPT) | IU/L | 15 – 50 | Sample Date | 2 | 95 | 13.00 | < 0.001 |
| ALT(SGPT) | IU/L | 15 – 50 | Days outside | 2 | 95 | 0.86 | 0.425 |
| ALP (Norm) | IU/L | 63 – 263 | Sample Date | 2 | 95 | 109.37 | < 0.001 |
| ALP (Norm) | IU/L | 63 – 263 | Days outside | 2 | 95 | 1.30 | 0.277 |
| GGT | IU/L | 2 – 51 | Sample Date | 2 | 95 | 5.37 | 0.006 |
| GGT | IU/L | 2 – 51 | Days outside | 2 | 95 | 0.01 | 0.987 |
| Total Bilirubin | mg/dL | 0.0 – 0.5 | Sample Date | 2 | 95 | 18.25 | < 0.001 |
| Total Bilirubin | mg/dL | 0.0 – 0.5 | Days outside | 2 | 95 | 1.03 | 0.363 |
| Creatinine | mg/dL | 0.6 – 1.6 | Sample Date | 2 | 95 | 3.83 | 0.025 |
| **Creatinine** | **mg/dL** | **0.6 – 1.6** | **Days outside** | **2** | **95** | **2.92** | **0.059** |
| BUN | mg/dL | 5 – 24 | Sample Date | 2 | 95 | 17.23 | < 0.001 |
| BUN | mg/dL | 5 – 24 | Days outside | 2 | 95 | 1.03 | 0.363 |
| BUN/Creatinine | -- | -- | Sample Date | 2 | 95 | 18.38 | < 0.001 |
| **BUN/Creatinine** | **--** | **--** | **Days outside** | **2** | **95** | **3.95** | **0.023** |
| Phosphorus | mg/dL | 4.0 – 9.0 | Sample Date | 2 | 95 | 2.42 | 0.094 |
| **Phosphorus** | **mg/dL** | **4.0 – 9.0** | **Days outside** | **2** | **95** | **3.60** | **0.031** |
| Glucose (Norm) | mg/dL | 50 – 90 | Sample Date | 2 | 95 | 31.23 | < 0.001 |
| Glucose (Norm) | mg/dL | 50 – 90 | Days outside | 2 | 95 | 1.69 | 0.190 |
| Ca (Norm) | mg/dL | 8.5 – 11.5 | Sample Date | 2 | 95 | 140.51 | < 0.001 |
| Ca (Norm) | mg/dL | 8.5 – 11.5 | Days outside | 2 | 95 | 1.39 | 0.255 |
| Mg (Norm) | mEq/L | 2.2 – 3.4 | Sample Date | 2 | 95 | 107.95 | < 0.001 |
| Mg (Norm) | mEq/L | 2.2 – 3.4 | Days outside | 2 | 95 | 0.48 | 0.620 |
| Na | mEq/L | 138 – 154 | Sample Date | 2 | 95 | 2.96 | 0.057 |
| Na | mEq/L | 138 – 154 | Days outside | 2 | 95 | 0.10 | 0.902 |
| K(Norm) | mEq/L | 3.4 – 5.5 | Sample Date | 2 | 95 | 12.91 | < 0.001 |
| K(Norm) | mEq/L | 3.4 – 5.5 | Days outside | 2 | 95 | 1.76 | 0.177 |
| Na/K (Norm) | -- | 30 – 42 | Sample Date | 2 | 95 | 18.21 | < 0.001 |
| Na/K (Norm) | -- | 30 – 42 | Days outside` | 2 | 95 | 0.86 | 0.425 |
| Chloride | mEq/L | 91 – 119 | Sample Date | 2 | 95 | 45.97 | < 0.001 |
| Chloride | mEq/L | 91 – 119 | Days outside | 2 | 95 | 0.50 | 0.608 |
| Cholesterol | mg/dL | 80 – 130 | Sample Date | 2 | 95 | 10.36 | < 0.001 |
| Cholesterol | mg/dL | 80 – 130 | Days outside | 2 | 95 | 0.46 | 0.635 |
| Triglyceride(Norm) | mg/dL | 20 – 75 | Sample Date | 2 | 95 | 9.95 | < 0.001 |
| **Triglyceride(Norm)** | **mg/dL** | **20 – 75** | **Days outside** | **2** | **95** | **6.50** | **0.002** |
| Amylase(Norm) | IU/L | -- | Sample Date | 2 | 95 | 3.30 | 0.041 |
| Amylase(Norm) | IU/L | -- | Days outside | 2 | 95 | 0.19 | 0.825 |
| Precision PSL | IU/L | -- | Sample Date | 2 | 95 | 62.96 | < 0.001 |
| Precision PSL | IU/L | -- | Days outside | 2 | 95 | 1.14 | 0.323 |
| CPK (Norm) | IU/L | 115 – 315 | Sample Date | 2` | 95 | 1.18 | 0.311 |
| CPK (Norm) | IU/L | 115 – 315 | Days outside | 2 | 95 | 1.05 | 0.356 |
| RBC | 10<sup>6</sup>/μL | 8.0 – 18.0 | Sample Date | 2 | 95 | 41.24 | < 0.001 |
| RBC | 10<sup>6</sup>/μL | 8.0 – 18.0 | Days outside | 2 | 95 | 1.21 | 0.302 |
| HGB | g/dL | 8.0 – 14.0 | Sample Date | 2 | 95 | 13.09 | < 0.001 |
| HGB | g/dL | 8.0 – 14.0 | Days outside | 2 | 95 | 1.95 | 0.150 |
| HCT | % | 10 – 39 | Sample Date | 2 | 95 | 30.71 | < 0.001 |
| **HCT** | **%** | **10 – 39** | **Days outside** | **2** | **95** | **3.02** | **0.054** |
| MCV | fL | 15 – 30 | Sample Date | 2 | 95 | 16.22 | < 0.001 |
| MCV | fL | 15 – 30 | Days outside | 2 | 95 | 1.07 | 0.347 |
| MCHC (Norm) | Pg | 35 – 42 | Sample Date | 2 | 95 | 23.08 | < 0.001 |
| MCHC (Norm) | Pg | 35 – 42 | Days outside | 2 | 95 | 1.93 | 0.151 |
| Platelet | 10<sup>3</sup>/μL | 120 – 550 | Sample Date | 2 | 95 | 4.72 | 0.011 |
| Platelet | 10<sup>3</sup>/μL | 120 – 550 | Days outside | 2 | 95 | 0.28 | 0.756 |
| Neutrophil | /μL | 1800 – 4500 | Sample Date | 2 | 95 | 23.16 | < 0.001 |
| Neutrophil | /μL | 1800 – 4500 | Days outside | 2 | 95 | 0.13 | 0.875 |
| Lymphocytes | /μL | 2500 – 7500 | Sample Date | 2 | 95 | 9.00 | < 0.001 |
| **Lymphocytes** | **/μL** | **2500 – 7500** | **Days outside** | **2** | **95** | **3.02** | **0.053** |
| Monocytes | /μL | 50 – 800 | Sample Date | 2 | 95 | 0.55 | 0.579 |
| Monocytes | /μL | 50 – 800 | Days outside | 2 | 95 | 0.33 | 0.722 |
| Eosinophils(Norm) | /μL | 0 – 800 | Sample Date | 2 | 95 | 2.16 | 0.120 |
| Eosinophils(Norm) | /μL | 0 – 800 | Days outside | 2 | 95 | 0.03 | 0.970 |
| Basophils(Norm) | /μL | 0 – 200 | Sample Date | 2 | 95 | 6.51 | 0.002 |
| Basophils(Norm) | /μL | 0 - 200 | Days outside | 2 | 95 | 1.76 | 0.178 |
| Bicarbonate | mEq/L | 20 – 25 | Sample Date | 2 | 95 | 91.95 | < 0.001 |
| Bicarbonate | mEq/L | 20 – 25 | Days outside | 2 | 95 | 0.620 | 0.539 |

All statistical procedures were performed using MiniTab 13.0 (Harrisburgh, PA). Statistical analysis examined the response of each hematological parameter to the number of days the goat had been in the study, the number of days it had spent on pasture, and the worm treatment it had undergone. These data were tested for normality using the Anderson-Darling Normality Test. If data were non-normal, they were transformed using the normalize function in MiniTab. Afterwards, data were analyzed using a General Linear Model to deduce if these timescales were predictive of the metric’s value. Then, multiple linear regression was run with all hematological parameters as predictors of “days on pasture” to express a relationship comparable to symptoms/signs of parasite load. Variance Inflation Factors (VIF) and the Predicted Residual Sums of Squares (PRESS) were evaluated. Afterwards, best-subsets regression was applied. Mallow’s statistic (c-p), the regression coefficient (*r^2^*) and the standard error of the regression (s) were evaluated to select the best available model. Multiple regression was run on this new model and the process of evaluating the VIF and PRESS repeated. If these values were not acceptable based on standard convention, we removed inflated predictors and ran the best-subsets regression again with the new selection of predictors. This was repeated until the best model was identified based on low VIF, PRESS, s, c-p, and high *r^2^*.

## Results

Six of the 38 hematological parameters screened were significantly increased as days of exposure to Coccidia increased (P = 0.001 – 0.031; Table 2). Additionally, Creatinine (P = 0.059), hematocrit (HCT; P = 0.054) and lymphocyte count (P = 0.053) increased with Coccidia exposure, but the relationship was marginally significant (Table 2).

Twenty-seven (71.1%) of the 38 hematological parameters made very significant changes over the duration of the study (P < 0.01; Table 2). Five showed significant changes (P = 0.01 – 0.041), and change in Sodium was marginally significant (P = 0.057) over the duration of the study.

The initial regression model contained the nine hematological predictors that had significant (P < 0.05) or marginally significant (0.05 < P < 0.1) relationships from the general linear models earlier reported herein. This model had relatively high VIF for Total Protein, Globulin, and Albumin (Table 3). Globulin’s VIF (48.4) was excessively high, and because globulin is also encompassed by total protein, it was removed from the regression. This brought VIF for all predictors to acceptable levels (Table 3).

**Table 3.**
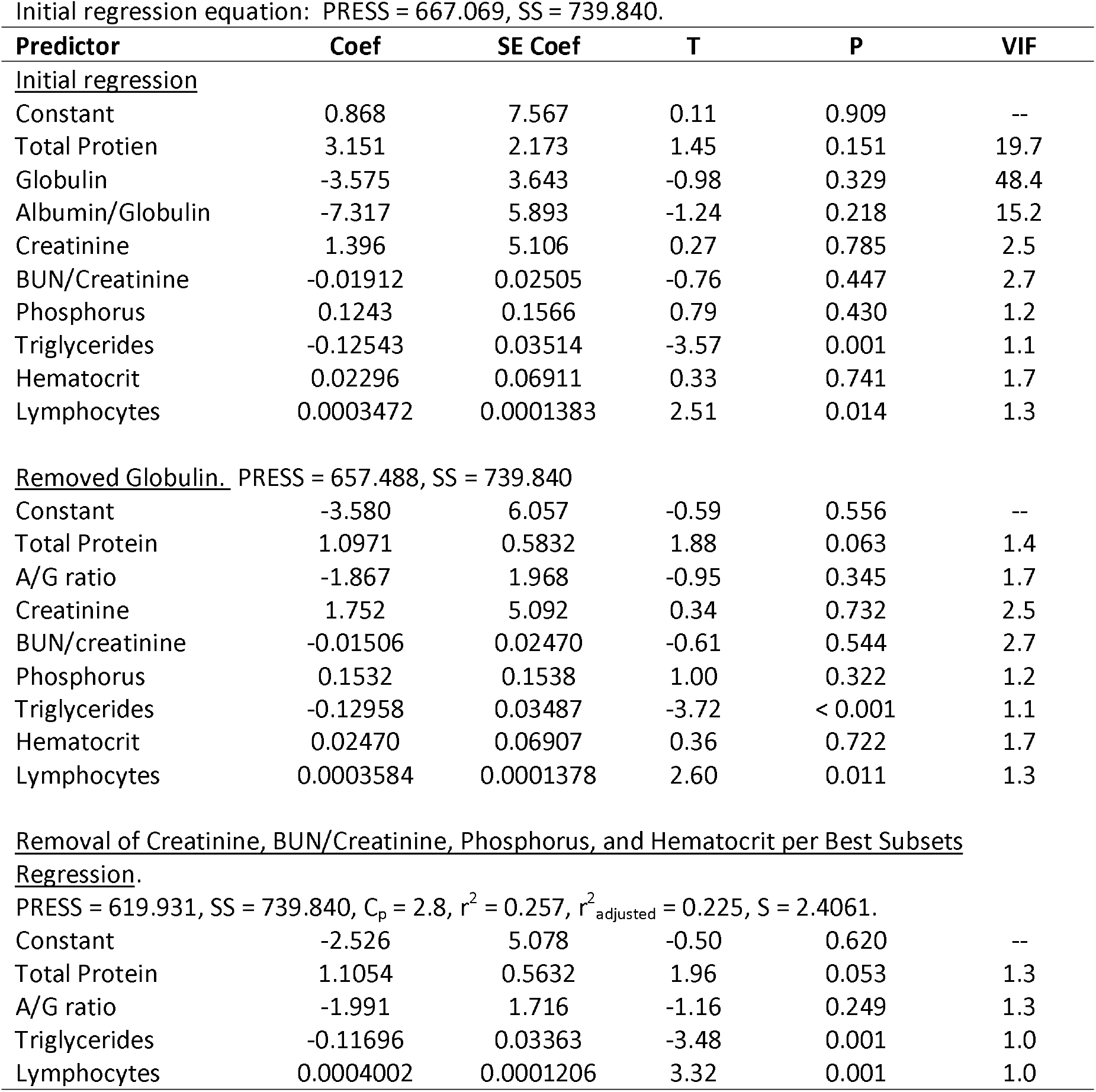
Linear regression modelling to predict exposure (days outside) to Coccidia.

Initial regression equation: PRESS = 667.069, SS = 739.840.
| Predictor | Coef | SE Coef | T | P | VIF |
| --- | --- | --- | --- | --- | --- |
| <u>Initial regression</u> |  |  |  |  |  |
| Constant | 0.868 | 7.567 | 0.11 | 0.909 | -- |
| Total Protien | 3.151 | 2.173 | 1.45 | 0.151 | 19.7 |
| Globulin | -3.575 | 3.643 | -0.98 | 0.329 | 48.4 |
| Albumin/Globulin | -7.317 | 5.893 | -1.24 | 0.218 | 15.2 |
| Creatinine | 1.396 | 5.106 | 0.27 | 0.785 | 2.5 |
| BUN/Creatinine | -0.01912 | 0.02505 | -0.76 | 0.447 | 2.7 |
| Phosphorus | 0.1243 | 0.1566 | 0.79 | 0.430 | 1.2 |
| Triglycerides | -0.12543 | 0.03514 | -3.57 | 0.001 | 1.1 |
| Hematocrit | 0.02296 | 0.06911 | 0.33 | 0.741 | 1.7 |
| Lymphocytes | 0.0003472 | 0.0001383 | 2.51 | 0.014 | 1.3 |
| Removed Globulin. PRESS = 657.488, SS = 739.840 |  |  |  |  |  |
| Constant | -3.580 | 6.057 | -0.59 | 0.556 | -- |
| Total Protein | 1.0971 | 0.5832 | 1.88 | 0.063 | 1.4 |
| A/G ratio | -1.867 | 1.968 | -0.95 | 0.345 | 1.7 |
| Creatinine | 1.752 | 5.092 | 0.34 | 0.732 | 2.5 |
| BUN/creatinine | -0.01506 | 0.02470 | -0.61 | 0.544 | 2.7 |
| Phosphorus | 0.1532 | 0.1538 | 1.00 | 0.322 | 1.2 |
| Triglycerides | -0.12958 | 0.03487 | -3.72 | < 0.001 | 1.1 |
| Hematocrit | 0.02470 | 0.06907 | 0.36 | 0.722 | 1.7 |
| Lymphocytes | 0.0003584 | 0.0001378 | 2.60 | 0.011 | 1.3 |
| Constant | -2.526 | 5.078 | -0.50 | 0.620 | -- |
| Total Protein | 1.1054 | 0.5632 | 1.96 | 0.053 | 1.3 |
| A/G ratio | -1.991 | 1.716 | -1.16 | 0.249 | 1.3 |
| Triglycerides | -0.11696 | 0.03363 | -3.48 | 0.001 | 1.0 |
| Lymphocytes | 0.0004002 | 0.0001206 | 3.32 | 0.001 | 1.0 |

The remaining eight predictors were subjected to Best Subsets Regression to evaluate the best models. This produced 36 possible models involving 1 – 8 predictors. We selected the following model

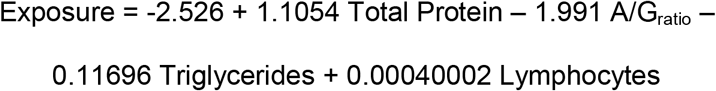

It was selected because it had the highest r^2^ (0.257) among all models with four or fewer predictors and the highest r^2^_adjusted_ (0.225) among all possible models providing it is the most precise model. Further, it had the strongest fit (C_p_ = 2.8) the best accuracy (s = 2.4061) among all possible models. There was no evidence that the levels of exposure to H.c. had any clinical influence on these numbers.

## Discussion

Our results supported observations by other researchers (opt. cite) that demonstrated physiological responses to coccidian infestation in vertebrate hosts and demonstrated that selected hematological parameters appear to have a relationship that can be modelled to predict levels of exposure and potentially infestation by coccidian in Alpine Dairy Goats, and the 100 year rain event stimulated infestation.

Previous studies of *Eimeria* infestation on hematology in goats exist. A controlled study of *Eimeria arloingi* investigated 10 hematological responses in 18 kid goats (Hashemnia et al. 2014). All of their experimental subjects demonstrated clinical symptoms of coccidiosis (e.g., diarrhea, oocyst excretion). Appearance of diarrhea was associated with reduced serum levels of Alkaline Phosphatase (ALP) activity, increased packed cell volume (PCV) and serum hemoglobin (HGB), and decreases in Sodium (Na^+^), Chloride (Cl^-)^, and Potassium (K^+^) levels. They found no treatment effects on serum levels of Aspartate Aminotransferase (AST), Alanine Aminotransferase (ALT), Gamma Glutamyltransferase (GGT), albumin, or total protein. Our report is ad hoc, so we did not sufficiently quantify oocysts or monitor diarrhea to evaluate these relationships. However, we did anecdotally observe occasional incidences of diarrhea among subjects. ALP activity did not change relative to the days on pasture; however, changes in PCV, HGB, Na^+^, Cl^-^, and K^+^ appeared to be symptomatic of confinement, not exposure to coccidia (Table 2). Like coccidia dosage in the above study, exposure via days on pasture did not result in altered AST, ALT, albumin, or GGT. However, each of these did change over the duration of our study (Fig. 1; Fig. 2), suggesting a confinement response or developing response to *H.c*. exposure that had yet to reach statistical significance.

**Figure 1.**
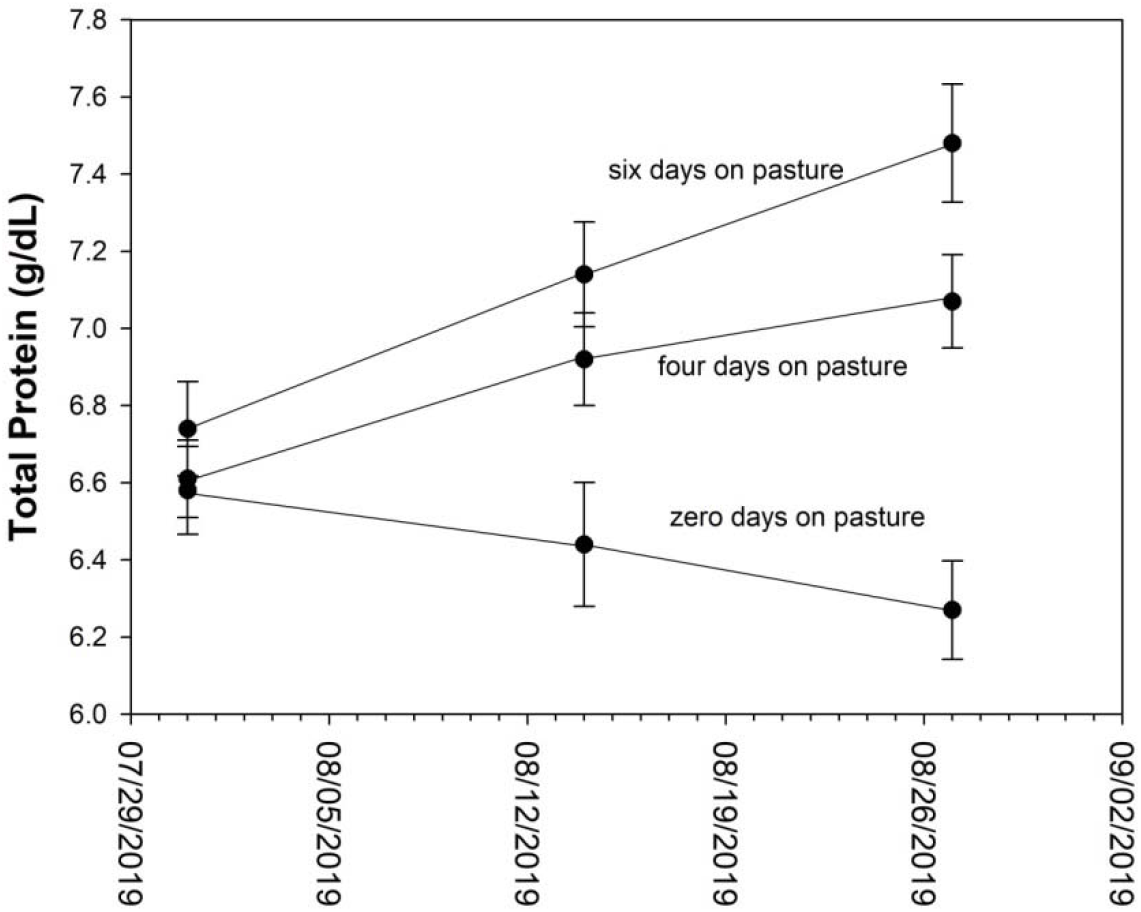
Physiological responses of Alpine Milk Goats to coccidian infestation during a 100-yr rain even in Oklahoma. Figure 1a. Response of Total Protein to time on pasture.

We observed total protein responses to time on pasture, supporting other studies with increased TP (El Manyawe et al. 2010; Singh et al. 2016; Mohamaden etal. 2018) and conflicting with the previously discussed study (Hasemnia et al. 2014). Whereas total protein fell among goats that had never been on pasture, it rose in those that spent time on it (Fig. 1a). If this response was due to high-quality pasture availability, it should have declined as the study ensured. Because it rose in goats that had been on pasture, but declined in those that had never been on pasture, it seems likely that the rise in total protein is a response to coccidian infestation. Increased total protein expression in response to coccidian infection is also known in chickens (Hirani et al. 2007; Koynarski et al. 2007; Gilbert eta l. 2011), sheep (Carzarotto et al. 2018), although reductions are also noted in sheep (Ghanem and Abd El-Raof 2005; Mohamaden et al. 2018). Generally, these relationships are complex because the age and previous exposure effects on total protein due to albumin and globulin effects (Fitzgerald 1964).

Pasture exposure also influenced globulin, A/G ratio, creatinine, BUN/creatinine ratio, phosphorus, triglycerides and hematocrit (Table 2). The increase in globulin among pastured goats compared to no significant change among unpastured goats suggests humoral response to the coccidian (Fig. 1b), as has been previously reported in goats (El Manyawe et al. 2010; Gwaze et al. 2010), and sheep (Carzarotto et al. 2018). Changes in albumin-globulin (A/G) ratio followed an inverted non-monotonic response curve over time and this variation was largely due to changes in globulin.

**Fig. 1b.**
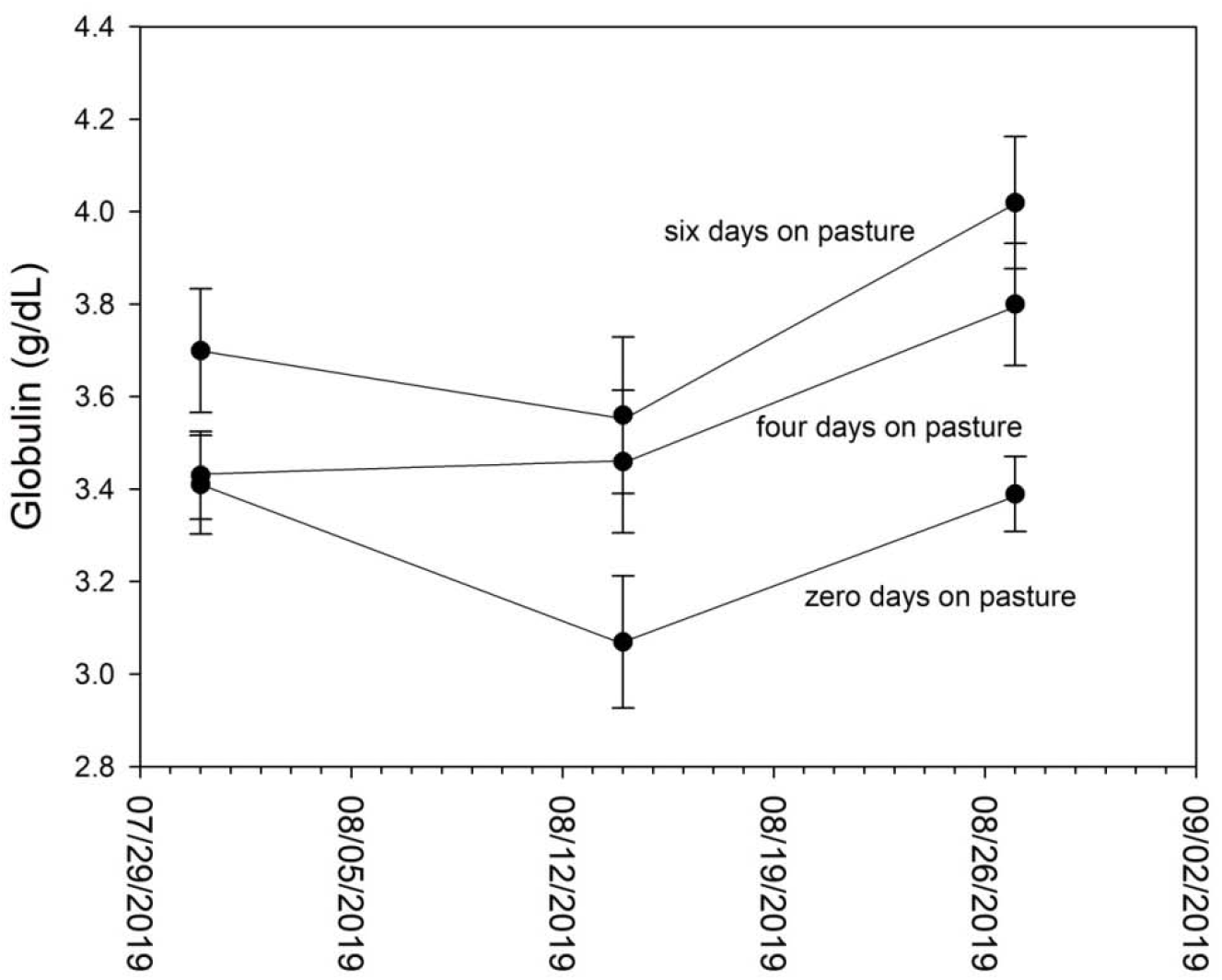
Globulin responses to days on pasture

Creatinine was also elevated in exposed goats (Fig. 2c.), as previously demonstrated (El Manyawe et al. 2010; Gwaze et al. 2010). This commonly signifies reduced renal clearance but arises in response to many kinds of stressors. Like A/G ratio, blood-urea-nitrogen (BUN)/creatinine ratio was reflective of the changes in creatinine while BUN remained unchanged. BUN is commonly a sign of dehydration, and both BUN and Creatinine rise during kidney disease. Since water was freely available, and we had no other evidence of kidney problems, this suggests that the coccidian assault and any possible associated secondary infections led to the rise in Creatinine.

Reduced triglyceride levels (Fig. 2g) may be indicative of disrupted fat absorption from rupture of luminal membranes as coccidian emerge from host cells. In fact, unpastured goats demonstrated much higher triglyceride levels than the pastured ones. There is some evidence of anticoccidial activity by medium-chain fatty acids; however, if goats increase production of these, our analysis is unable to discern this against losses from intestinal injury.

Hematocrit fell for all groups during the duration of the study, however, pastured goats had somewhat higher percentages than unpastured goats. Reduced hematocrit is expected in animals exposed to coccidian because of the blood loss associated with sloughing luminal membranes (da Silva et al. 2012). However, it is also associated with nutritional problems including low iron, B12, and folate. These animals were fed a balanced diet; however, malabsorption of these nutrients due to coccidian-induced damage is also a reasonable expectation.

As we observed, serum lymphocyte numbers rise in response to coccidiosis in goats (Shommein and Osman 1980; Jenkins et al. 1991). Leukocytosis involving T-cell proliferation is known from chickens that were previously immunized (Jenkins et al. 1991). Further, mobilization of lymphocytes (mostly B-cells) from Peyers patches is also common (Gregory 1990). However, leukocytopenia may also arise (Jenkins et al. 1991).

The predictive model produced in this study demonstrates that such models for clinical usage using patterns of multiple indicators of pathogenesis may be useful as for earlier diagnosis coccidiosis in goats and other organisms. Improved methods of diagnosis of coccidiosis are critical to reducing incidences of coccidiosis and the economic efficiency of livestock operations (Silva et al. 2013; Chartier and Paraud 2012; Tauseef-ur-Rehman et al. 2011). Currently, diagnosis of coccidiosis is commonly dependent on screening feces for oocysts of the parasites involved (ConsTable 3020) although other methods do exist (Khodakaram-Tafti et al. 2017; Silva et al. 2013). However, our model is not a direct relationship to coccidian numbers, and is therefore not truly translatable for clinical use in its current form. It relays a surrogate, days on pasture, to opportunistically test proof-in-concept of a tool for this purpose. The current model depends on data collected in the wettest year in the recorded history of Oklahoma, thus it also provides evidence of the impacts of lengthy extreme weather expected under climate change. But, it is not representative of a typical year, thus a multi-year study is needed to construct a true predictive model for use. Further, different breeds may respond differently to coccidian infections, especially when comparing meat to dairy breeds. However, the model we produced provides evidence that this next step can be taken. As such, with refinement, ground-truthing, and validation, a usable model would be a useful tool for practioners. In this way, a simple blood test plugged into a spreadsheet equation would assist diagnosis and its need for monitoring or treatment before animals become symptomatic, thus improving health management, biosecurity, and medical costs to a producer. Consequently, future studies should focus on translating this information from days of exposure to actual parasite loads, and comparing a diversity of breeds to identify differences is needed before this concept will be available for clinical diagnosis.

Climate change is an important topic at this time. Our results arise from a 100 year rain event that is representative of what the future holds. Major rain events such as this can threaten the goat industry by raising parasite control costs and reducing production efficiency. It is important that the goat industry proactively consider management approaches to deal with these coming challenges before they become commonplace.

**Fig. 1c.**
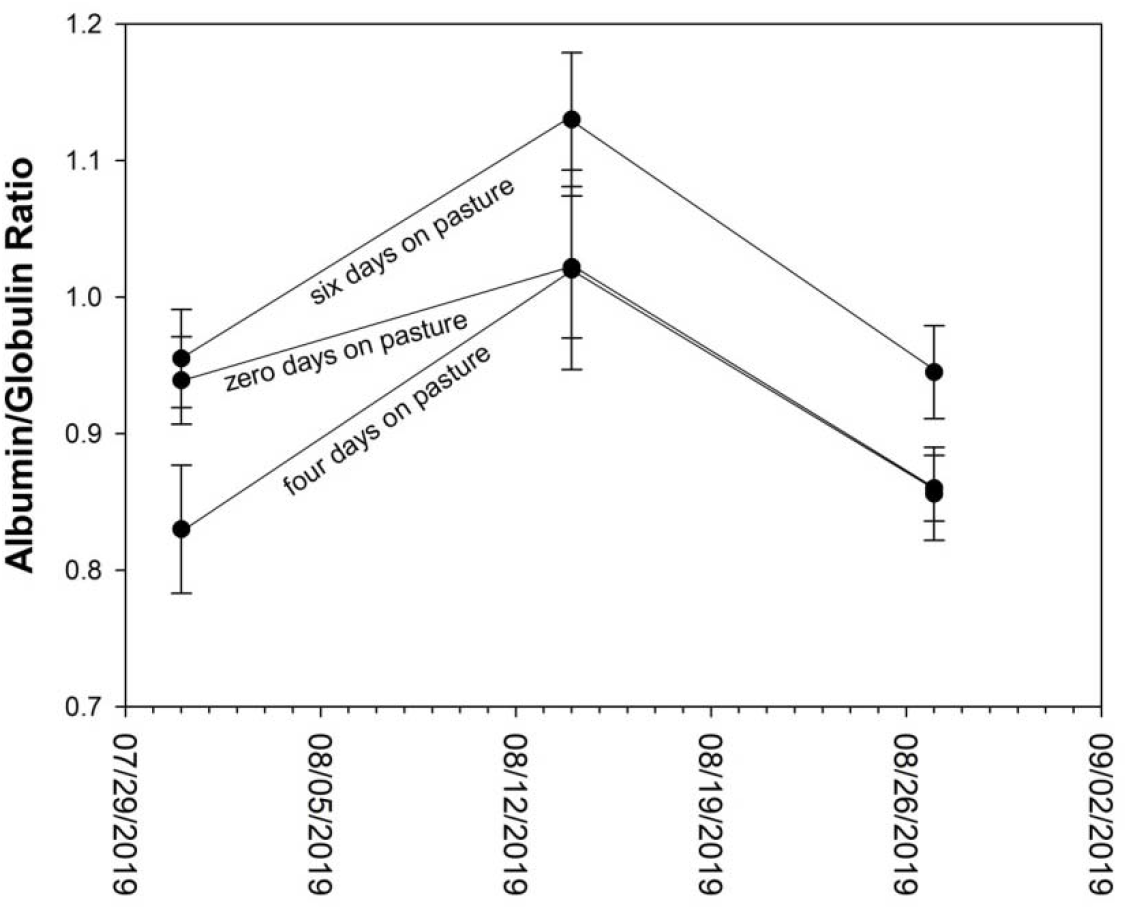
Albumin/Globulin Ratio.

**Fig. 1d.**
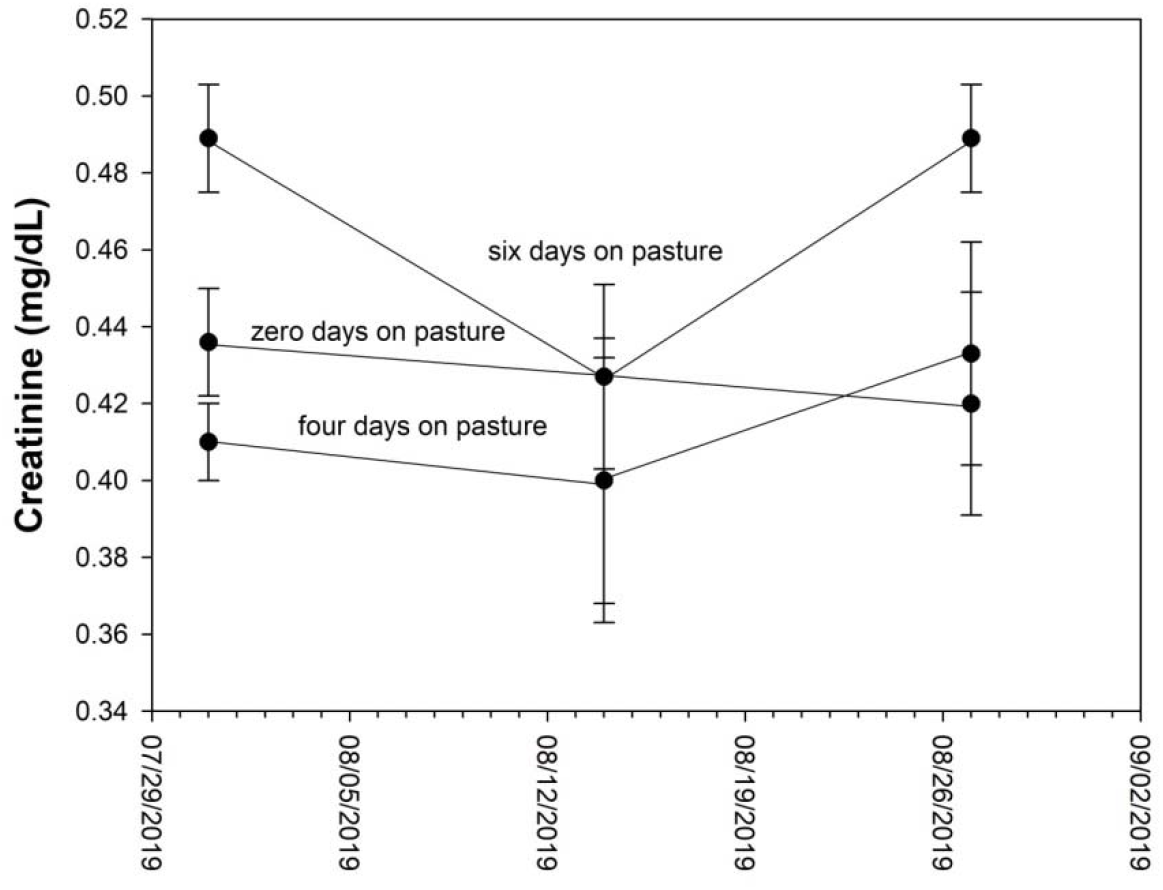
Creatinine

**Fig. 1e.**
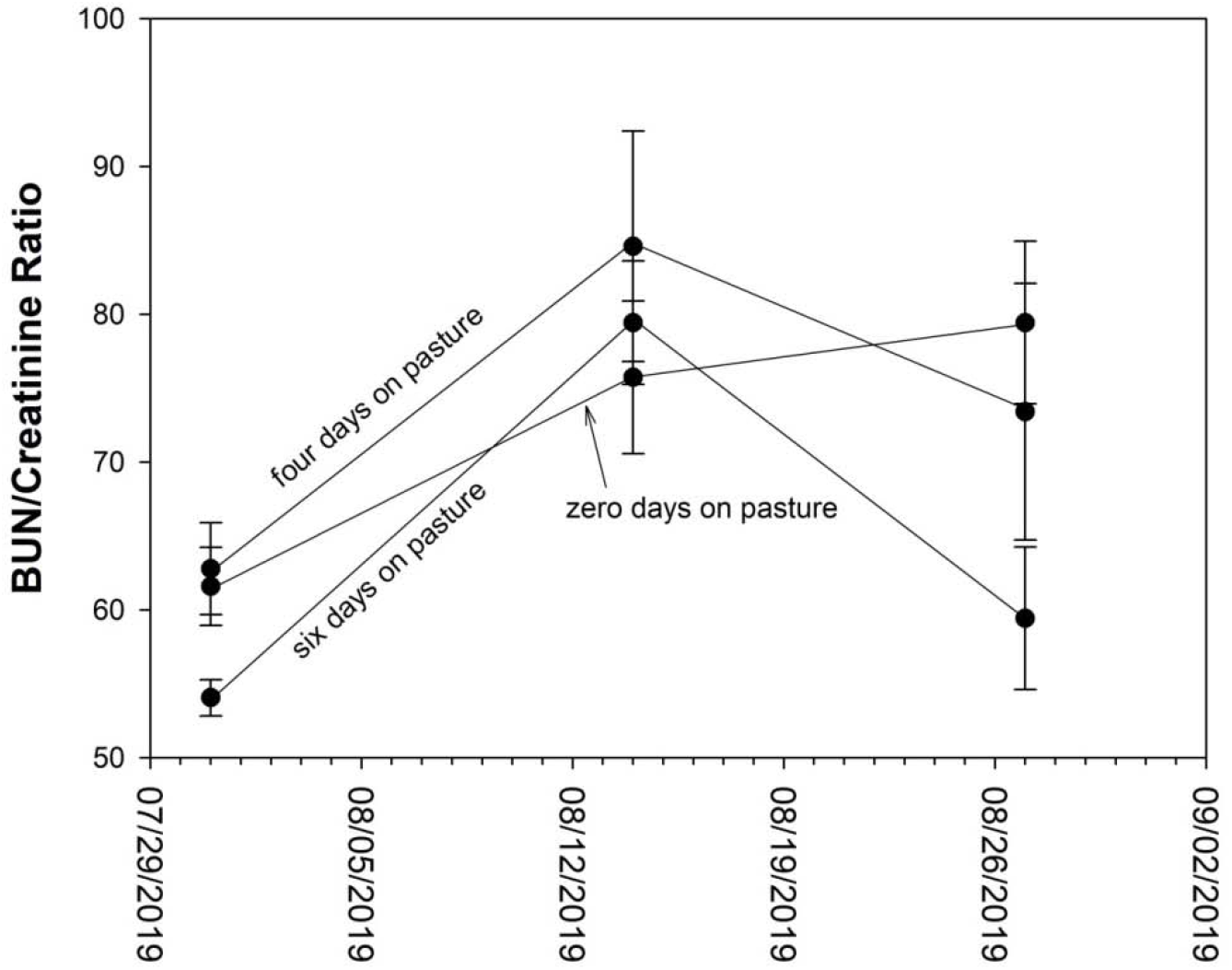
BUN/Creatinine Ratio.

**Fig. 1f.**
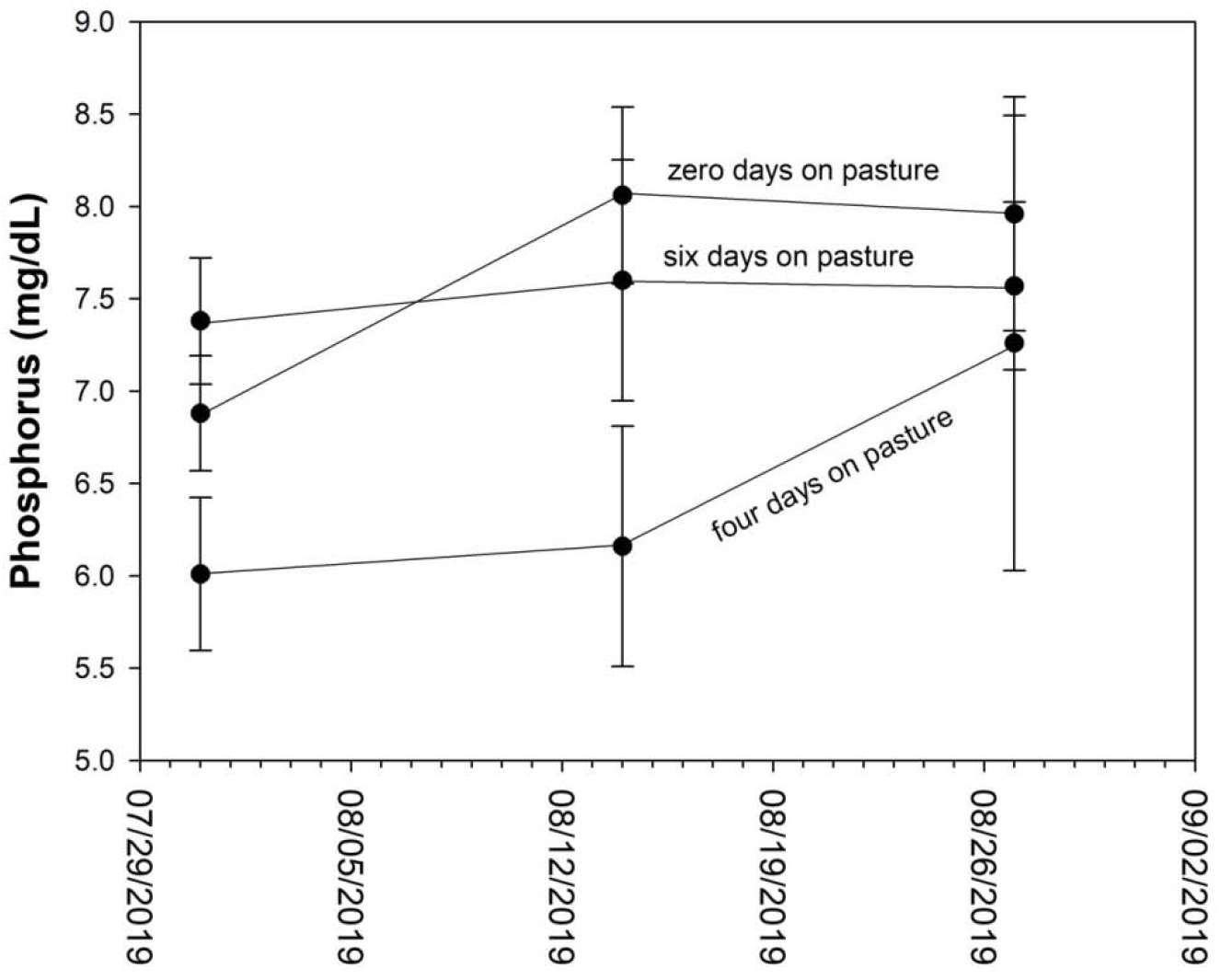
Phosphorus.

**Fig. 1g.**
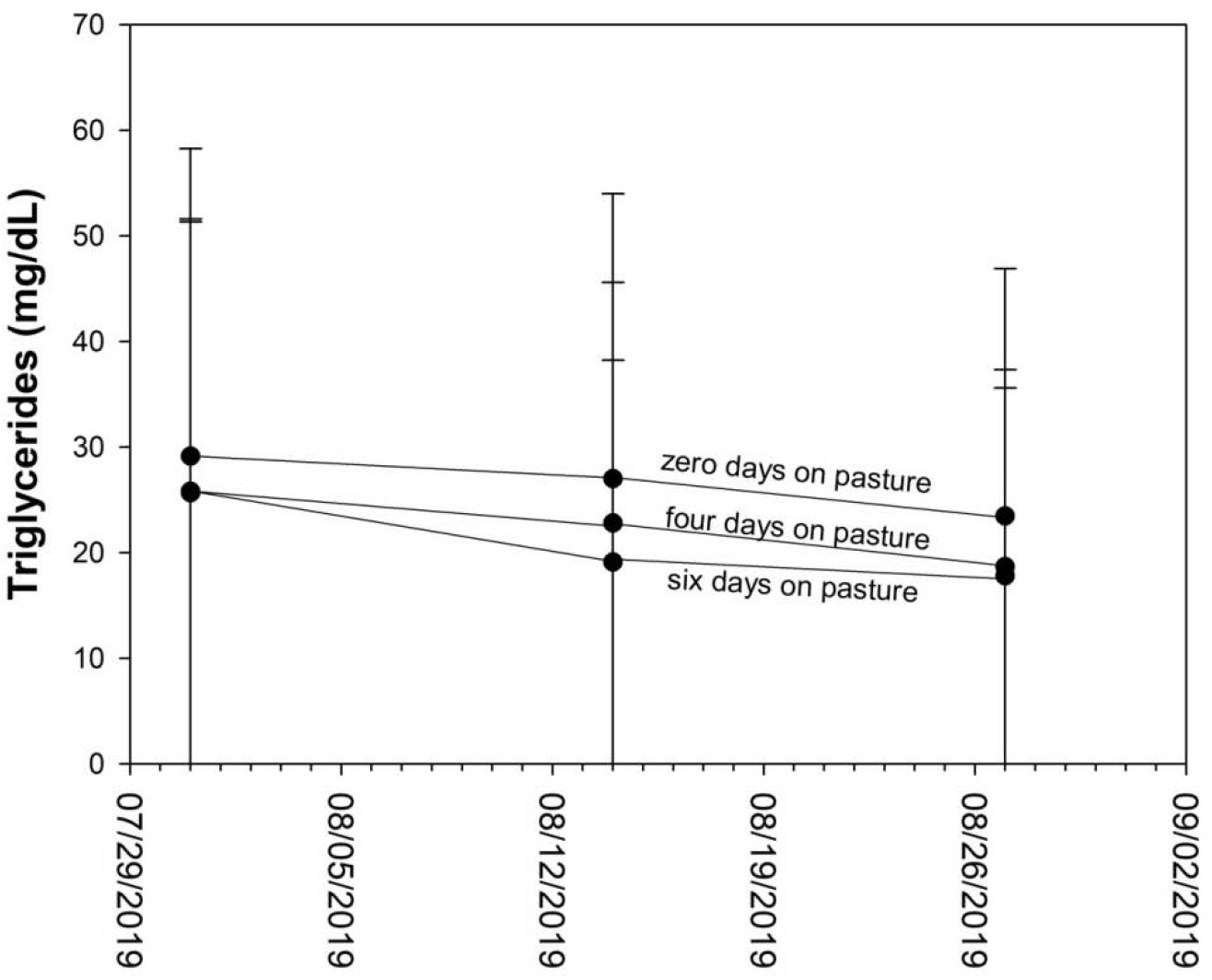
Triglycerides.

**Fig. 1h.**
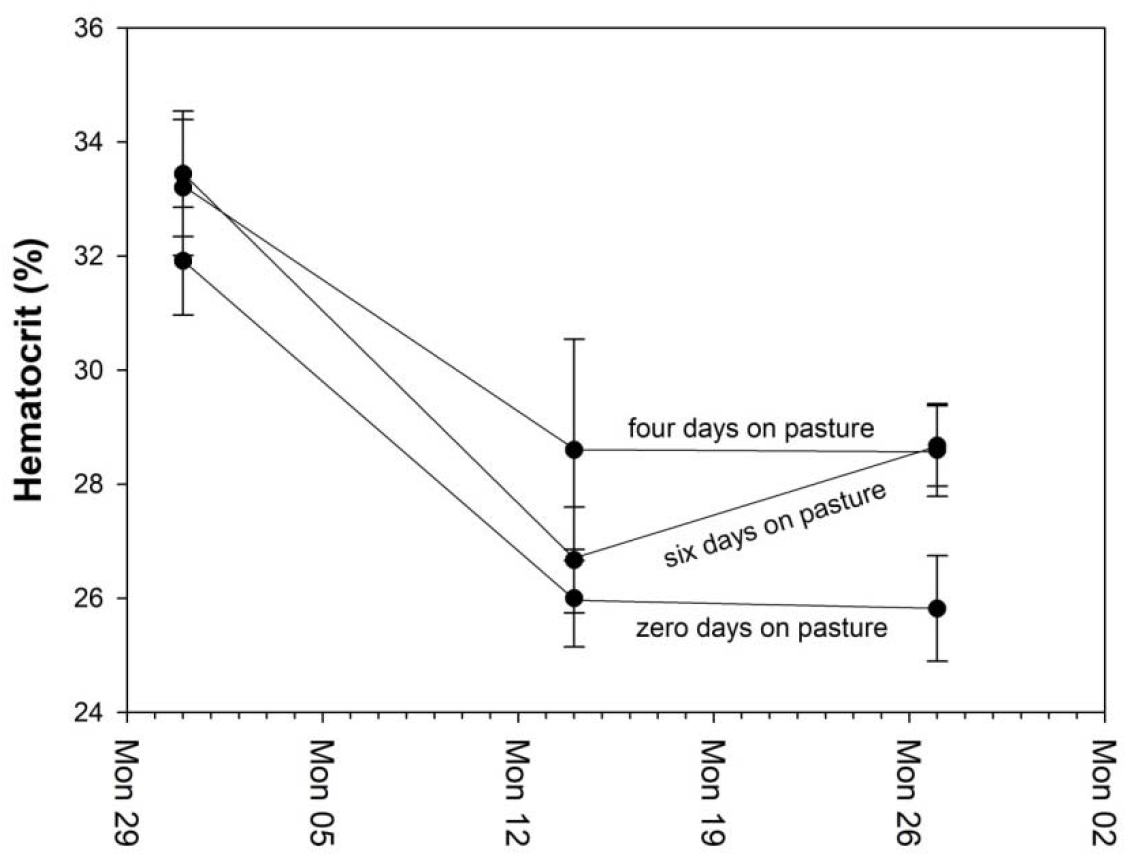
Hematocrit

**Fig. 1i.**
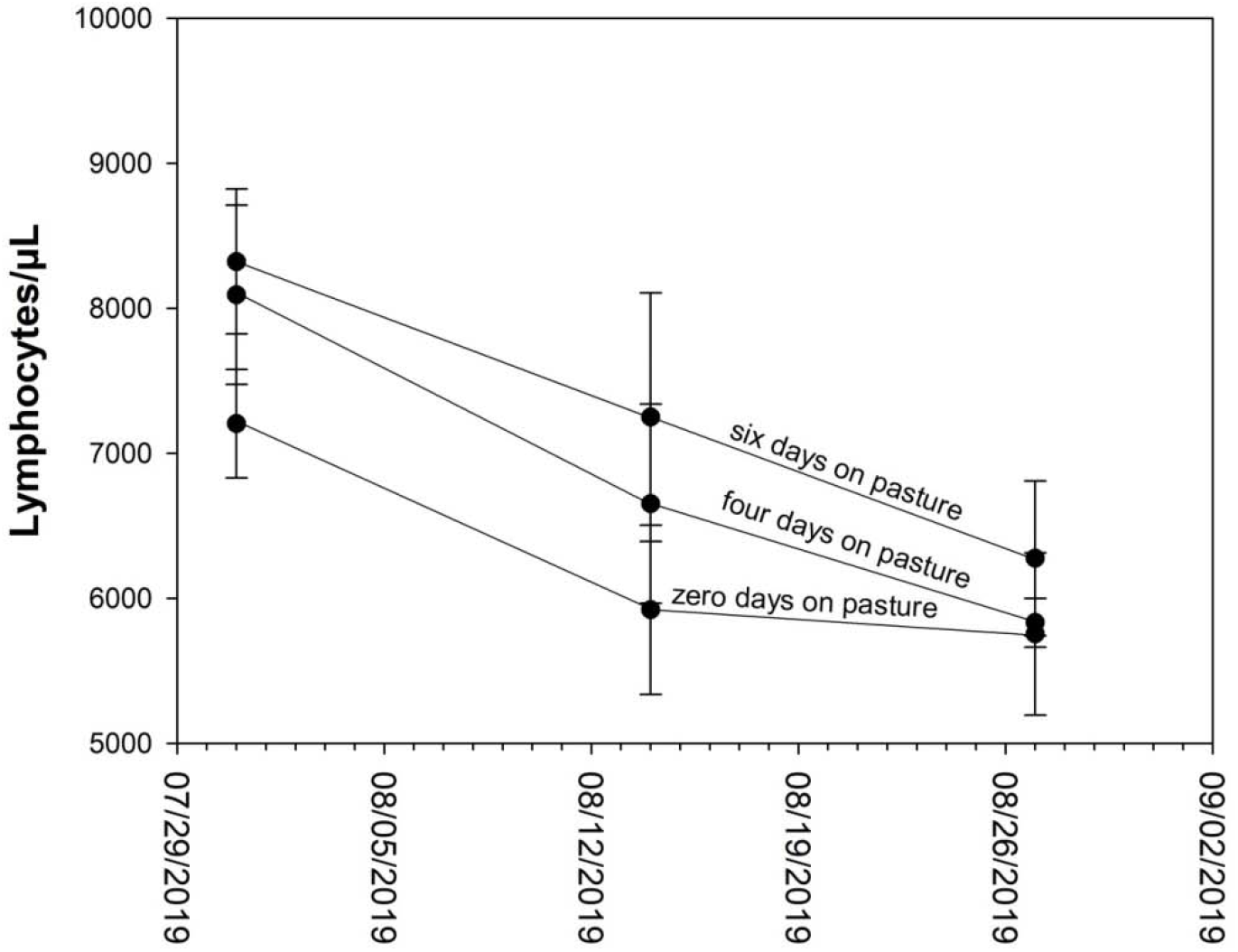
Lymphocytes.

## Acknowledgements

This study was funded by USDA NIFA Evans-Allen funds (Project # OKLUTILAHUN2018) and protocols were approved under the Animal Welfare Act by the Langston University Institutional Animal Care and Use Committed (Approval #2018-14).

